# Productivity of *Pleurotus ostreatus* on lignocellulosic Rice Husk substrate supplemented with Sugar Industry Residual Biomass

**DOI:** 10.1101/2025.03.17.643815

**Authors:** Erwin Manuel Hernández García, Jhonatan Jhosseppy Romero Morales, Marilín Nicoll Sánchez Purihuamán, Ada Patricia Barturen Quispe, Lizbeth Maribel Córdova Rojas, Sebastian Huangal Scheineder, Segundo Alberto Vasquez Llanos, Carmen Rosa Carreño Farfán

## Abstract

Substrate formulations based on *Oryza sativa* (rice) husk supplemented with sugar industry waste bagasse and *Saccharum officinarum L*. filter cake were used as an alternative substrate to cultivate mushrooms such as *Pleurotus osteatrus*, being essential the comparison of the productivity parameters carried out as carpophores produced per productive cycle, productive period, biological efficiency and production rate, in addition to the physicochemical analysis of the lignocellulosic substrates, which in the case of organic matter, nitrogen, phosphorus and potassium are in the ranges of 28.37-35.20%; 1.28-1.45%; 0.06-1.74%; 0.46-1.17%, respectively. Regarding productivity parameters, carpophores with a total weight of 83, 99.67 y 101 g were harvested in a ratio of rice husks (CA) with sugar cane bagasse (BC) of 66.7:33.3, 50:50 and 25: 75, respectively. In the case of biological efficiency, it was 27.66-50.50 % with rice husk plus sugarcane bagasse and 8.67-9.51 % with rice husk plus filter cake compared to 8.56 % in the rice husk control and the production rate was 0.58-0.92. Based on these findings, it was found that the addition of sugarcane bagasse significantly favoured the productivity of *Pleurotus ostreatus* in the rice husk, obtaining not only a higher yield but also a higher biological efficiency, in addition to improving the nutritional quality of the fungi, where the highest protein production was obtained in rice husks supplemented with sugarcane bagasse in the 25: 75 proportion.

**RESUMO:** Formulações de substratos à base de casca de “arroz” de *Oryza sativa* suplementada com bagaço de resíduo da indústria sucroalcooleira e torta de filtro de *Saccharum officinarum L*. foram utilizadas como substrato alternativo para o cultivo de cogumelos como *Pleurotus Osteatrus*, sendo essencial a comparação dos parâmetros de produtividade realizados como carpóforos produzidos por ciclo produtivo, período produtivo, eficiência biológica e taxa de produção, além da análise físico-química dos substratos lignocelulósicos que no caso da matéria orgânica, nitrogênio, fósforo e potássio estão nas faixas de 28,37-35,20%; 1,28-1,45%; 0,06-1,74%; 0,46-1,17%, respectivamente. Em relação aos parâmetros de produtividade, foram colhidos carpóforos com peso total de 83, 99,67 e 101 g nas proporções de casca de arroz (CA) e bagaço de cana (BC) de 66,7:33,3, 50:50 e 25:75, respectivamente. No caso da eficiência biológica, foi de 27,66 - 50,50% com casca de arroz mais bagaço de cana e 8,67 - 9,51% com casca de arroz mais torta de filtro, em comparação com 8,56% na casca de arroz controle e a taxa de produção foi de 0,58 - 0,92. Com base nesses achados, verificou-se que a adição de bagaço de cana-de-açúcar favoreceu significativamente a produtividade de *Pleurotus ostreatus* na casca de arroz, obtendo não só maior rendimento como também maior eficiência biológica, além de melhorar a qualidade nutricional dos fungos, onde a maior produção de proteína foi obtida na casca de arroz suplementada com bagaço de cana-de-açúcar na proporção 25:75.

*Palavras-chave:* Eficiência biológica, proteína, biomassa residual.

## 1. INTRODUCTION

Every year around the world, 1.3 billion of food or agro-industrial waste are produced (Tlais et al., 2020), Global food demand is significantly influenced by population growth and per capita income growth, food demand is projected to increase by up to 102% by 2050, especially in developing countries (Fukase et al., 2020). It is estimated that food waste contributes to the emission of around 3.3 million tons of CO2 into the atmosphere annually. (Capanoglu et al., 2022), this is why lignocellulosic waste from agro-industrial activity has gained importance because it is an alternative for obtaining new economically and environmentally valuable products, highlighting the importance of the transition towards a circular and sustainable bioeconomy. (Blasi et al., 2023)

Worldwide, around 540 million tons of sugarcane bagasse are generated as a by-product of the sugar industry, similarly, the “rice husk” of *Oryza sativa*, which is a by-product of rice milling, generates approximately 130 million tons of rice husk per year (Awasthi et al., 2024). The incineration of this waste introduces pollutants into the air that negatively affect the environment and human health. (Wang et al., 2011). In June 2024 in Peru, the generation of rice husk was estimated at 1.9 million tons (SIEA., 2024); an option for the use of this waste is through the cultivation of edible mushrooms, including P*leurotus ostratus*, where lignocellulosic waste such as rice husks and sugarcane bagasse products can be used (Akcay et al., 2023) taking into account that for each ton of ground stems, 27-28% bagasse is produced (Bezerra & Ragauskas, 2016), 3 % of sugarcane filter cake and ash (Minh et al., 2023).

Rice husk contains 35-40% cellulose, 15-20% hemicellulose and 20-25% lignin (Gao et al., 2018), so it can be used for the cultivation of edible mushrooms (Ríos et al., 2017) such as *Pleurotus* ostreatus, which have enzymes such as versatile peroxidase that degrade these substrates (Andlar et al., 2018). Likewise, it is possible that rice husk and sugarcane bagasse, may have greater biological efficiency when processed with other substrates to optimize nutrient availability (Doroški et al., 2022).

The research addressed one of the problems of the rice and sugar cane industry, which is the poor use of by-products and the agro-industrial processing generates husk, bagasse and sugarcane filter cake, which are mostly not used optimally to give them the added value they should have. Edible mushrooms of the *Pleurotus* ostreatus genus have a protein conversion efficiency per unit of area and time superior to animals. This fungal genus is the second most commercialized worldwide after *Agaricus* spp. and has high yield and productivity; the fruiting bodies are less susceptible to attack by insect pests and phytopathogens and are recognized for their medicinal properties. The objective of this study is to determine the productivity of *Pleurotus ostreatus* with a lignocellulosic substrate of rice husk (*Oryza sativa*) supplemented with residual biomass from the sugar industry, aiming to validate a methodology that can be replicated in different places, with the assurance of obtaining a safe and quality product for human consumption.

## 2. MATERIALS AND METHODS

*Pleurotus ostreatus* seeds were inoculated in the investigated substrates and incubated for 15-18 days; fruiting or basidiocarp production was induced for 10-15 days and harvesting was carried out in two waves (inoculation, incubation, fruiting, and harvesting. In the explanatory research, the wholly randomized experimental design (DCA) was used with nine treatments, corresponding to T1: Control 100% rice husk (RH); T2: Control 100% sugarcane bagasse (SB); T3: 100% Control sugarcane filter cake (SFC); T4: 66.7% RH + 33.3% SB; T5: 50% RH + 50% SB; T6: 25% RH + 75% SB; T7: 66.7% RH + 33.3% SFC; T8: 50% RH + 50% SFC and T9: 25% RH + 75% SFC. Three replicates were performed in each treatment, with a total of 27 experimental units. Agro-industrial companies in the Lambayeque Region provided the biomasses under study.

### 2.1 Characterization of rice husks, bagasse and sugarcane filter cake

The substrates rice husk, bagasse and sugar cane filter cake were analyzed in the Pro Suelos y Aguas (LABSAF) laboratory of the National Institute of Agrarian Innovation (INIA) in Chiclayo, where the pH, electrical conductivity, organic matter, nitrogen, phosphorus, potassium, calcium, magnesium, dry matter, humidity, ash, carbon and carbon-nitrogen ratio were determined.

### 2.2. Obtaining the mother strain from *Pleurotus ostreatus*

The carpophores of *Pleurotus ostreatus* were purchased from the Biotechnology Company of Fungi Innova - Lima and were taken to the laboratory of the Research Center for Sustainable Development (CIFOS) for the isolation of the secondary mycelium. The carpophores were washed in drinking water, disinfected with 1% sodium hypochlorite for 1 minute, rinsed eight consecutive times with distilled water and dried with sterilized filter paper. Next, triangular fragments (1 cm on each side) (Figure 2A, 2B) were cut from the fruiting body with a scalpel, placed on Petri dishes with glucose potato agar (PGA) plus chloramphenicol (six fragments per dish) and incubated at 28 °C for 10 - 15 days. Once the secondary mycelium had developed, blocks of agar containing the mycelium (1 cm^2^) were cut, placed in tubes with inclined PDA and incubated at 28 °C until the mycelium colonized the culture medium. This way, the secondary mycelium or “mother strains” of *Pleurotus ostreatus* were obtained and stored in a refrigerator at eight °C. (Mendoza et al., 2019).

### 2.3. Obtaining seeds from *Pleurotus ostreatus*

The wheat grains with the husk were selected to remove the damaged ones, washed with tap water five consecutive times, placed in a pot with enough water to double their volume and cooked until the grains were in a “chalky” state (15 minutes). The grains were then placed in a sieve to remove the remaining water, spread out on a solid surface for 30 minutes to reduce excess moisture and mixed with 4.5 g of calcium carbonate and 17.5 kg of gypsum/kg of wheat. The grains were placed in wide-mouthed glass jars, occupying 2/3 of the content, covered with aluminum foil and paper, sterilized in an autoclave at 15 pounds of pressure, 21°C for 1 hour, and cooled for 12 hours before sowing the fungus (Angulo Zubieta et al., 2022).

The mother strain of *Pleurotus ostreatus* was plated on PDA plates (1 cm^2^ fragments) and incubated at 30°C for 10 days. A block of PDA agar (2 x 2 cm) was placed on the surface of the sterilized wheat grains, and on top of this, a block of *Pleurotus ostreatus* mycelium with the agar included. Incubation was carried out at 30°C for 10 days until the mycelium thoroughly colonized the wheat grains (Mendoza et al., 2019)

### 2.4. Conditioning of substrates for the cultivation of *Pleurotus ostreatus*

The sugarcane filter cake and bagasse were dried at room temperature (22 – 24°C) for 15 days. The sugar cane filter cake was then crushed and sieved to remove unwanted material. The substrates, rice husk, bagasse, and sugarcane filter cake, were weighed and placed in polyethylene bags (8 x 12 cm) according to the corresponding treatments, and the required weight was determined based on the volume needed to fill ¾ of the capacity of the polypropylene bags. The rice husk was soaked in potable water for 48 hours at a rate of 1 liter of water per 100 g of husk and was drained with a cloth to remove excess water. Once the substrate mixtures were prepared, they were placed in polypropylene bags (8 x 12 cm) previously perforated with holes (diameter of 5 mm), weighed, closed with adhesive tape and pasteurized at 80 °C for 60 minutes. Afterward, the substrates were drained at room temperature (22 – 24 °C) for 48 hours until reaching 65 – 75 % moisture (Cruz, 2021).

### 2.5. Inoculation and incubation of substrates

The previously pasteurised and drained substrates were inoculated with 20% of the seed (60, 90 and 120 g depending on the bag weight). For inoculation, the substrate was placed on a previously disinfected aluminium tray, carefully mixed with the seed, and then the inoculated substrate was packed in polypropylene bags (8 x 12 cm) with a hermetic seal. A vertical cut (3 cm) was made on the central part of both sides of the bags with the help of a scalpel to facilitate aeration. The substrates were incubated on a closed wooden shelf (dark) with a temperature range of 22 - 25 ° C for the mycelium to fully colonize the substrate. The observation of humidity on the walls of the bags indicated the beginning and growth process of the fungus. The days required to colonize 50 and 100% of the substrates by the secondary mycelium of *Pleurotus ostreatus* were recorded (Nieto Juárez et al., 2021).

### 2.6. Production of carpophores or fruiting

The bags with the substrates entirely colonised by the mycelium of *Pleurotus ostreatus* were taken to a ventilated environment (window 116 x 136 cm), average temperature 18.9 °C; average relative humidity of 85 ± 5 %; illumination for 12 hours a day (fluorescent tubes at a light intensity of 500 lux units) and were sprayed with running water with a dispenser every 12 hours. In this environment, the production of primordia, the formation of closed carpophores, and the opening of carpophores were observed for 3 days (Nieto Juárez et al., 2021).

### 2.7 Harvesting and fruiting

The harvest was carried out using the “torsion” method, 4 days after the opening of the carpophores, for which the stem of fruiting bodies or “pineapple” was taken by its base and with a slight downward movement it was extracted. Next, the height and weight of the pineapple were determined, as well as the number, diameter and weight of each carpophore. The carpophores were sent to the Microservilab E.I.R.L laboratory to determine the protein content in percentage using the Kjeldahl method. During fruiting, the following parameters were recorded: time (days) of colonization of 50% of the substrates by secondary mycelium; for fruiting the following parameters were recorded: time (days) of colonization of 100% of the substrates by secondary mycelium, time (days) from sowing to the first harvest, weight of “pineapple” of fresh carpophores per bag per harvest, number and diameter of carpophores per pineapple, time (days) from sowing to the second harvest, weight of fungi per production cycle (first and second harvest) and protein content of the carpophores (Figure 2C, 2D) (Mendoza et al., 2019).

### 2.8. Comparison of productivity parameters of *Pleurotus osteatrus*

The productivity of *Pleurotus ostreatus* was expressed in terms of grams of fresh carpophores per production cycle, productive period PP (days) (Akter et al., 2022), biological efficiency and production rate in the rice husk mixed of the lignocellulosic substrate with bagasse and sugarcane filter cake: fresh carpophores per production cycle (g), productive period, PP (days).

Biological efficiency (BE) was calculated according to the following equation (Equation 1): (De Mastro et al., 2023)

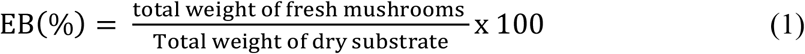

Likewise, the Production Rate (TP) (D. D. C. D. M. ruby O. A. benitez Cruz, 2021): (Equation 2):

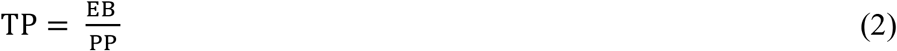

Where in: PP – productive period, days.

## 3. RESULTS AND DISCUSSION

### 3.1. Physical characteristics of the raw material

The lignocellulosic substrates rice husk, bagasse and sugarcane filter cake presented 28.37 - 35.20 % of organic matter, 1.28 - 1.45 % of nitrogen, 0.06 - 1.74 % of phosphorus, and 0.46 - 1.17 % of potassium. The highest values of organic matter, nitrogen, and phosphorus corresponded to sugarcane filter cake and potassium in rice husks. Regarding pH, it was slightly acidic (rice husk), acidic (sugarcane filter cake), and very acidic (sugarcane bagasse). The salinity level (CEc) was low for the three substrates (1.59 - 7.13 mhos/cm), and the humidity was 10.05 % (husk), 39.15 % (bagasse), and 47.40 % in filter cake. Regarding the carbon/nitrogen ratio, it ranged from 12.85 to 14.08. The three residues presented organic matter, carbon and nitrogen, better performance, nutrients, and a condition required to cultivate edible mushrooms such as *Pleurotus ostreatus* (Akcay et al., 2023; Desisa et al., 2023; Tang et al., 2019). Rice husk contains cellulose that varies between 25 and 45%, hemicellulose between 11 and 20%, and lignin between 20 and 35.33% (Wu et al., 2018). In sugarcane bagasse, cellulose has been reported to vary between 30-40%, hemicellulose 19-28%, lignin 17-25% and ash 5-20% (Antunes et al., 2022; Melesse et al., 2022).

Among the physical and chemical characteristics of the substrates used (Table 1) to cultivate *Pleurotus ostreatus*, pH and carbon-nitrogen ratio are the most important. The pH should be 5 – 6.5 (Zubairi et al., 2022). Lower values inhibit the mycelial development of the fungus. The carbon-nitrogen ratio shows the carbon content used for the growth of the mycelium and the formation of fruiting bodies or carpophores. Concerning texture, it also influences the development of the fungus because it is related to the diffusion of oxygen. More time was required for mycelial colonization in substrates with sugarcane filter cake in their composition (Desisa et al., 2024).

**Table 1.** Physical and chemical characteristics of the husk of *Oryza sativa L*., bagasse and sugarcane filter cake of *Saccharum officinarum L*.

| Physicochemical characteristics | *Oryza sativa* L husk. | Sugarcane bagasse | Sugarcane filter cake |
| --- | --- | --- | --- |
| pH | 6,70 | 4,00 | 5,40 |
| CEc (mmhos/cm) | 2,26 | 1,59 | 7,13 |
| Organic matter (%) | 28,37 | 32,15 | 35,20 |
| Nitrogen (%) | 1,28 | 1,37 | 1,45 |
| Phosphorus (P<sub>2</sub>O<sub>5</sub>) (%) | 0,06 | 0,16 | 1,74 |
| Potassium (K<sub>2</sub>O) (%) | 1,17 | 1,05 | 0,46 |
| Calcium (CaO) (%) | 1,57 | 1,98 | 3,52 |
| Magnesium (MgO) (%) | 0,60 | 0,83 | 1,37 |
| Dry matter (%) | 89,95 | 60,85 | 52,60 |
| Moisture (%) | 10,05 | 39,15 | 47,40 |
| Ash (%) | 4,76 | 6,15 | 23,40 |
| Carbon (%) | 16,45 | 18,65 | 20,42 |
| Rate C/N | 12,85 | 13,61 | 14,08 |

\* Soil and Water Analysis Laboratory, National Institute of Agrarian Innovation (INIA)

The use of rice husk for the cultivation of *Pleurotus ostreatus* coincides with (Akcay et al., 2023) who also used other lignocellulosic residues such as hazelnut branches, hazelnut shells, wheat straw and coffee waste; (Quiñónez-Martínez et al., 2022) they used plant species *Solanum elaeagnifolium* and *Salsola kali*., (Munir et al., 2024) pineapple residues; (Lozano Rocha et al., 2023) brewer spent grain (Wakawa et al., 2023) con olive pruning residues and spent coffee grounds.

Cellulose degradation by *Pleurotus ostreatus* involves the cellulolytic system of endoglucanases, cellobiohydrolases, and β-glucosidases. Endoglucanases degrade glycosidic bonds within the cellulose molecule, cellobiohydrolases or exoglucanases release cellobiose units from the ends, and β-glucosidases hydrolyze cellobioses with the release of glucose as the final product (Díaz Muñoz et al., 2019). Lignin is degraded by the oxidative lignocellulolytic system consisting of peroxidases, manganese peroxidases and laccases that open the phenyl rings and cause unstable free radicals that polymerise (Daz Muoz et al., 2019; Pia-Guzmán et al., 2016).

The degradation of lignocellulosic substrates by *Pleurotus ostreatus* was quantified by (Rodriguez et al., 2018) who analysed the pear pomace after carpophores harvest and determined a significant decrease in organic matter, soluble carbohydrates and fibre fractions, as evidenced by the reduction of neutral detergent fibre corresponding to lignin, cellulose, and hemicellulose (NDF), acid detergent fibre (ADF), lignin in acid detergent (LDA), cellulose (CEL) and hemicellulose (HCEL).). For its part, Östbring et al. (2023) demonstrated that *Pleurotus ostreatus* obtained with the deproteinized rapeseed cake substrate are useful for mushroom production and as a food source.

### 3.2. Carpophores of *Pleurotus ostreatus* obtained in the lignocellulosic substrate of rice husk supplemented with bagasse and sugarcane filter cake

Regarding the carpophores of *Pleurotus ostreatus* obtained in the lignocellulosic substrate of rice husk supplemented with sugarcane bagasse and sugarcane filter cake, the mycelium of *Pleurotus ostreatus* colonized 100 % of the wheat grains after 12 days. It constituted the seed used for the inoculation of the investigated lignocellulosic substrates. The mycelium of *Pleurotus ostreatus* colonized 13.33 - 30.00 % (5 days); 33.33 - 53.33 % (10 days); 86.66 – 100 % (15 days) and 100 % (20 days) of the substrates consisting of rice husk with 33.3 % (T4), 50 % (T5) and 75 % (T6) of sugarcane bagasse. Likewise, the mycelium of *Pleurotus ostreatus* colonized 10.00 – 13.33 % (5 days); 23.33 % (10 days); 53.33 – 83.33 % (15 days) and 100 % (20 days) of the substrates consisting of rice husk with 33.3 % (T7), 50 % (T8) and 75 % (T9) of sugarcane filter cake compared to 3.33 – 13.33 % (5 days); 13.33 – 23.33 % (10 days); 13.33 – 66.66 % (15 days) and 16.16 – 100 % (20 days) in the corresponding controls. (Figure 1.)

**Figure 1.**
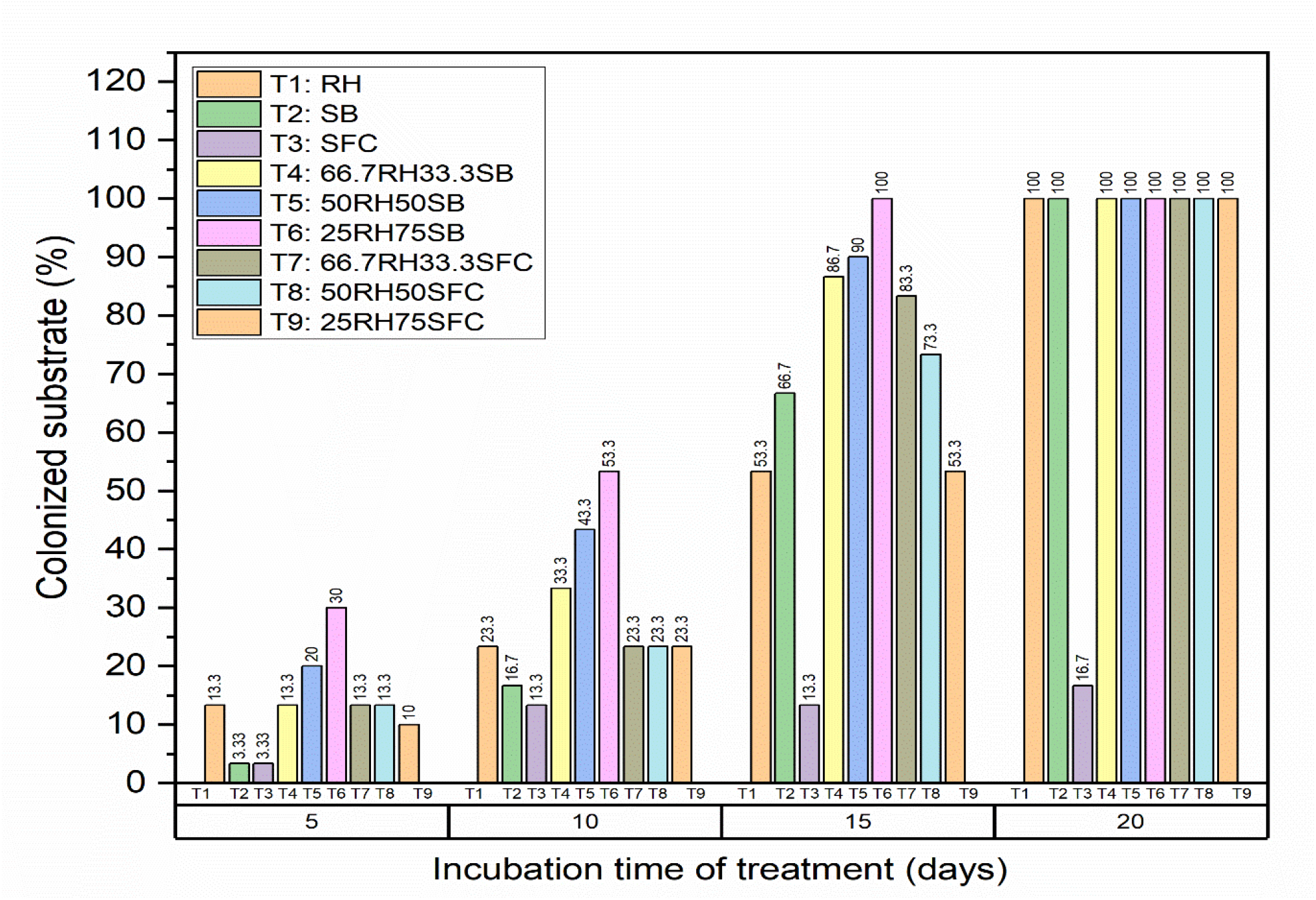
Percentage of lignocellulosic substrate colonized by *Pleurotus ostreatu*s mycelium at 5, 10, 15 and 20 days.

**Figure 2.**
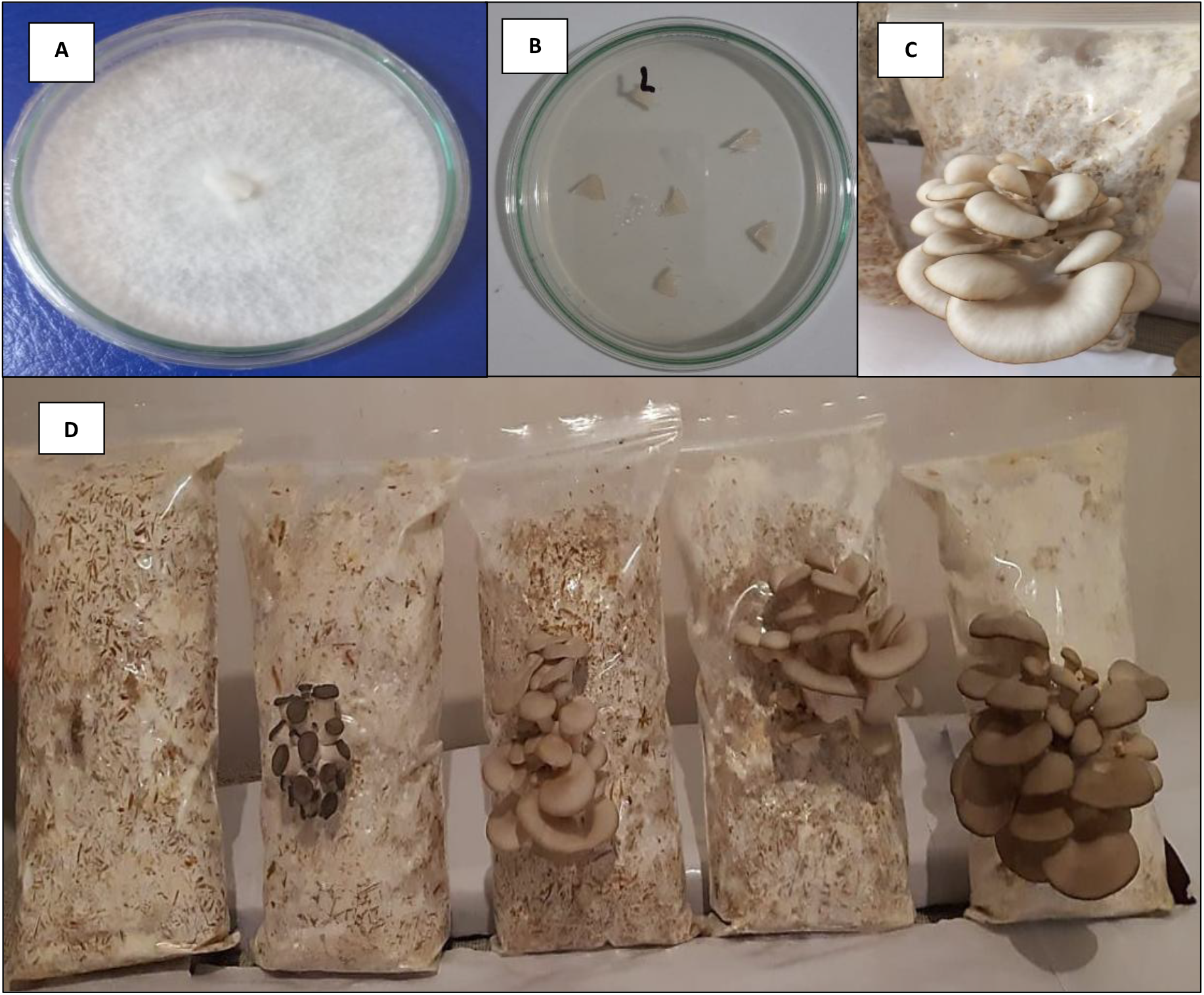
**A** Mycelium of *Pleurotus ostreatu*s grown on potato dextrose agar; **B:** *Pleurotus ostreatus* fragments cultured on Potato dextrose agar; **C**: *Pleurotus ostreatus* carpophores harvested from the cultivation of *Oryza sativa* husk with *Saccharum officinarum L*. bagasse; **D**: Sequence of formation and growth of *Pleurotus ostreatus* carpophores in *Oryza sativa* husk with *Saccharum officinarum L* bagasse (25% CA + 75% BC).

The time required for colonization by the *Pleurotus ostreatus* mycelium decreased as the percentage of sugarcane bagasse increased. On the contrary, the time increased as the percentage of sugarcane filter cake increased. In this context, 8-13 days were needed for 50% colonization and 15-20 days for 100% colonization of rice husk with sugarcane bagasse. In turn, 13 days were required for 50% colonization and 18-20 days for colonization of rice husk with filter cake (Figure 1)

Once 100% of the lignocellulosic substrates had been colonized (15-20 days), the primordia appeared after 22-23 days in the rice husk with sugarcane bagasse and after 25-41 days in the rice husk with sugarcane filter cake, compared to 29 days in the rice husk control. In the second harvest, primordia appeared after 42-49 days in the rice husk with sugarcane bagasse and after 44 days in the rice husk with sugarcane filter cake (Table 2). The primordia, which were initially white, grew, became greyish and produced carpophore, whose opening was observed after 3-5 days in the rice husk with sugarcane bagasse and after 3-5 days in the rice husk with sugarcane filter cake. In the second harvest, the carpophores opened after 2-3 days in the rice husk with sugarcane bagasse and did not open in the rice husk with sugarcane filter cake.

**Table 2.** Characteristics of *Pleurotus ostreatus* carpophores harvested on different lignocellulosic substrates.

| Treatments | First harvest |  |  |  | Second harvest |  |  |  | Total of carpophores |  |
| --- | --- | --- | --- | --- | --- | --- | --- | --- | --- | --- |
|  | Days to harvest | Weight of cone (g) | Number of carpophores | Carpophore diameter range (cm) | Days to harvest | Weight of cone (g) | Number of carpophores | Carpophore diameter range (cm) | Weight (g) | Number |
| T1 | 39 | 25.67 | 6.00 | 1 – 9 | 0 | 0 | 0 | 0 | 25.67 | 6.00 |
| T2 | 30 | 75.00 | 13.67 | 1 – 13 | 51 | 20.33 | 2.33 | 3 – 8 | 95.33 | 16.00 |
| T3 | 0 | 0 | 0 | 0 | 0 | 0 | 0 | 0 | 0 | 0 |
| T4 | 29 | 76.00 | 11.00 | 2 – 10 | 48 | 7.00 | 2.33 | 1 – 8 | 83.00 | 13.33 |
| T5 | 29 | 76.33 | 14.67 | 1 – 15 | 55 | 23.33 | 2.00 | 1 – 10 | 99.67 | 16.67 |
| T6 | 30 | 74.00 | 18.67 | 1 – 9 | 55 | 27.00 | 3.67 | 5 – 9 | 101.00 | 22.33 |
| T7 | 40 | 35.67 | 3.33 | 1 – 13 | 0 | 0 | 0 | 0 | 35.67 | 3.33 |
| T8 | 48 | 17.33 | 1.00 | 7 – 11 | 0 | 0 | 0 | 0 | 17.33 | 1.00 |
| T9 | 0 | 0 | 0 | 0 | 0 | 0 | 0 | 0 | 0 | 0 |

The carpophores were harvested at 29-30 days (first harvest) and 48-55 days (second harvest) after sowing the *Pleurotus ostreatus* seed in the rice husk with sugarcane bagasse. In turn, the first harvest was carried out at 40-48 days (first harvest) and no second harvest was observed in the rice husk with sugarcane filter cake. In the rice husk control, the first harvest was carried out in 39 days, and no second harvest was observed. The weight of the cone, the number of carpophores and the range of the diameter of the carpophores were more significant in the first than in the second harvest. The total weight of the carpophores was 83-101 g and the number was 13.33-22.33 in the rice husk with sugarcane bagasse. The total weight was 17.33 – 35.67 g and 1.00 – 3.33 in the rice husk with sugarcane filter cake. The rice husk control, the values were 25.67 and 6.00, respectively (Table 2)

### 3.3. Comparison of the productivity parameters of *P. ostreatus* in the lignocellulosic substrate rice husk supplemented with sugarcane bagasse and sugarcane filter cake

The mycelium of *Pleurotus ostreatus* colonized the sugarcane husk, bagasse and sugarcane filter cake; however, the sugarcane filter cake was only colonized up to 16% and primordia and carpophores were not formed, the appearance of primordia was at 4 days, the harvest at 167 days and the biological efficiency of 15.65% compared to the crushed bark with 21 days, 117 days and 48.20% respectively. Lignocellulosic substrates are suitable for cultivating *P. ostreatus*; however, to increase the biological efficiency and the production rate, mixtures are made with different substrates such as hazelnut branches, hazelnut shells, wheat straws, and spent coffee grounds (Akcay et al., 2023), cottonseed husk with sugarcane bagasse, chicken manure (Desisa et al., 2024), beet residues (Östbring et al., 2023).

The addition of sugarcane bagasse to rice husks favoured the cultivation of Pleurotus ostreatus, and the positive effect increased as the concentration of agroindustrial residue increased. In this context, a decrease in the number of days required for the colonization of the substrate by the mycelium and an increase in biological efficiency and production rate were observed, a result that can be explained by the increase in aeration in the substrate mixture. The cultivation of edible mushrooms is an aerobic process favoured by substrates that provide nitrogen (Vieira & de Andrade, 2016) and moisture retention of more than 65% (Wiesnerová et al., 2023)

The weight of the carpophores increased according to the percentage of sugarcane bagasse, 83 g with 33.3 % sugarcane bagasse; 99.67 g with 50 % sugarcane bagasse and 101 g with 75 % sugarcane bagasse. On the contrary, the weight of the carpophores decreased as the percentage of sugarcane filter cake increased: 35.67 g with 33.3 % sugarcane filter cake, 17.33 g with 50 % sugarcane filter cake, and no carpophores were harvested with 75 % sugarcane filter cake (Table 3). In both harvests, the weight of the carpophores was 25.67 g in the rice husk control. The time elapsed from the sowing of *Pleurotus ostreatus* mycelium to the last harvest was 39-55 days, and the average number of carpophores per production cycle was 13.33-22.33 in rice husk with sugarcane bagasse. In turn, the values were 48 days and 1.0-3.33 carpophores in rice husk with sugarcane bagasse (Table 2).

**Tabla 3.**
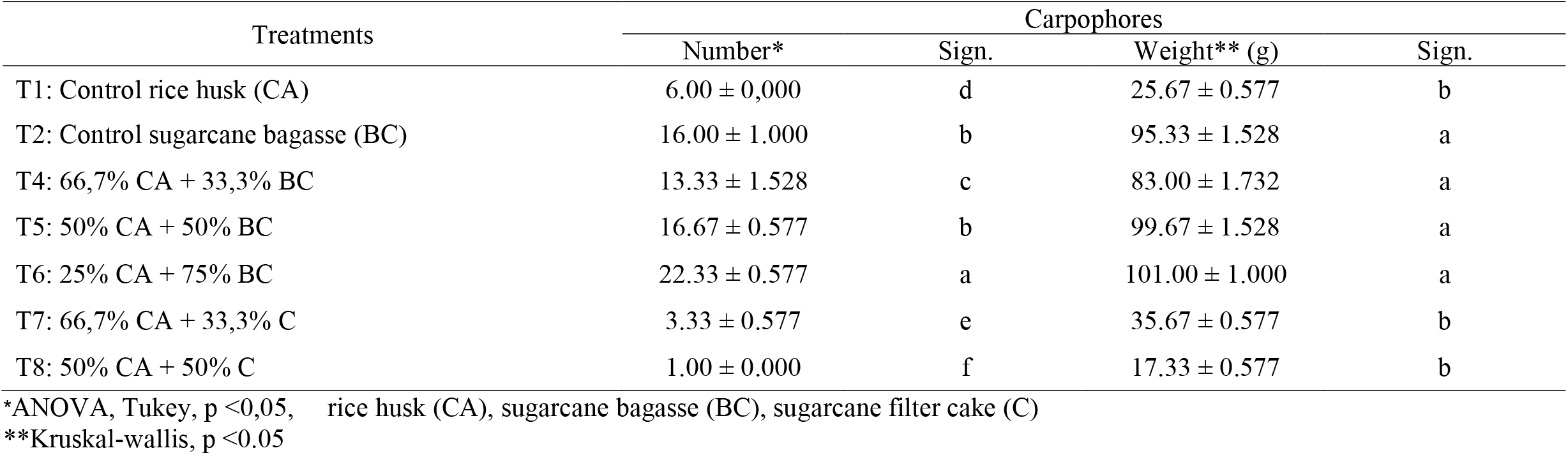
Multiple comparison test of the number and weight of carpophores of *Pleurotus ostreatus* grown on different lignocellulosic substrates.

For the rice husk control, the values were 39 days and 6 carpophores per production cycle. Statistically, the number of carpophores of *Pleurotus ostreatus* presented normality and homogeneity of variances, the parametric test analysis of variance demonstrated significance and the highest value (22.33) corresponded to T6, with significant differences with respect to the other treatments, statistically the values of the carpophores weight did not present normality, the nonparametric Kruskall-Wallis test demonstrated significance and the highest weights corresponded to T6, T5, T4 and T2 with significant differences with respect to T1, T7 and T8 (Table 3).

The 100% colonization of rice husk and the obtaining of carpophores demonstrated that it constitutes a substrate to cultivate *Pleurotus ostreatus*; however, the biological efficiency was 8.56% (Table 4). In this regard, Akcay et al. (2023) reported that rice husk alone had a biological efficiency of 45.6, while with spent coffee grounds in a 1:1 ratio, its performance was 64.3% and they concluded that the residue is effective only when mixed with another residue.

**Table 4.** Multiple comparison tests of *Pleurotus ostreatus* productivity parameters with different lignocellulosic substrates.

| Treatments | Weight of carpophores<br>per production cycle (g) | Productive period<br>(días) | Biological efficiency* (%) |  | Production rate* |  |
| --- | --- | --- | --- | --- | --- | --- |
|  |  |  | Valor | Sign. | Valor | Sign. |
| T1: Control rice husk (CA) | 25.67 | 39 | 8.56 ± 0.196 | c | 0.22 ± 0.006 | c |
| T2: Control sugarcane bagasse (BC) | 47.67 | 51 | 31.78 ± 0.508 | b | 0.62 ± 0.010 | b |
| T4: 66,7% CA + 33,3% BC | 41.50 | 48 | 27.66 ± 0.577 | b | 0.58 ± 0.012 | b |
| T5: 50% CA + 50% BC | 49.83 | 55 | 49.83 ± 0.764 | a | 0.91 ± 0.015 | a |
| T6: 25% CA + 75% BC | 50.50 | 55 | 50.50 ± 0.500 | a | 0.92 ± 0.010 | a |
| T7: 66,7% CA + 33,3% C | 35.67 | 40 | 9.51 ± 0.156 | c | 0.24 ± 0.006 | c |
| T8: 50% CA + 50% C | 17.33 | 48 | 8.67 ± 0.289 | c | 0.18 ± 0.006 | c |

\* Kruskal-wallis, p <0,05

The addition of sugarcane filter cake to rice husk negatively affected the biological efficiency and the production rate of *Pleurotus ostreatus* and the impact was more significant as the residue concentration increased, a result that can be explained by the higher density (0.25 g m-3) of the sugarcane filter cake compared to sugarcane bagasse (0.12 g m-3).). The higher the density, the greater the compaction, and therefore there is less aeration. Bagasse is a fibrous residue obtained after grinding sugarcane to extract the juice, producing approximately 27-28% of the dry weight of the juice. (Bezerra & Ragauskas, 2016). For its part, sugarcane filter cake is a by-product of sugar cane, a spongy and amorphous material that is greenish in colour and obtained by the action of temperature during the surface filtration of the juice for the production of panela. (Lagos-Burbano & Castro-Rincón, 2019).

The negative effect of substrate compaction during *Pleurotus ostreatus* cultivation is corroborated by (Josephat et al., 2020) who determined as the bran concentration increased (from 25 to 75%) mixed with eucalyptus bark (from 75 to 25%), the number of days required for mycelium colonization increased (from 20 to 29) and, on the contrary, the biological efficiency decreased from 32.95 to 13.87%. Likewise, (Puig-Fernández et al., 2020) They reported that the growth of *Pleurotus ostreatus* mycelium was not uniform in the brewer’s bran and hypothesised that the high content of fine particles caused compaction of the substrate, hindered the aeration required for the fungi, and promoted anaerobic conditions that caused the death of the fungal mycelium.

The productivity of mushroom cultivation is evaluated by the parameters of biological efficiency (BE) parameters expressed in kg of fresh mushroom biomass per kg of dry substrate used (Östbring et al., 2023), productive period (PP) (Zárate-Salazar et al., 2020) and the production rate (TP) (Akcay et al., 2023). The EB is used to evaluate the quality of an organic residue as a substrate to cultivate edible mushrooms and is classified as acceptable between 40 – 50%, the minimum reference value for the economically profitable commercial cultivation of *P. ostreatus* (Piña et al., 2017). The mixture of rice husk with sugarcane bagasse for *Pleurotus ostreatus* cultivation favoured EB, a result that coincided with Akcay et al. (2023) who achieved 64.3% in the treatment of sugarcane bagasse with spent coffee ground (1:1); (Bernarda et al., 2017) hey reached 177.37% of EB with 15% of rice husk, 40% of lentil stover, 40% of barley straw, 3% of soybean meal and 2% of corn stover (Table 4).

The number of harvests increased with the substrate mixture, which was two compared to the control, where one harvest was achieved or none. Regarding the productive period (PP) in the present investigation, it was 55 days in two harvest waves compared to the values reported by Rodríguez et al. (2018) with 45 days and (Zárate-Salazar et al., 2020) with 60 days after three waves. According to (Muswati et al., 2021), the higher the biological efficiency, the shorter the productive period. The mixture of rice husk with bagasse also favoured the production rate of *Pleurotus ostreatus*, as reported by (Mendoza et al., 2019), with a value of 0.75 with the mixtures of coffee plus white bolaina shavings, compared to the control in which 0.47 was reached. (bolaina shavings) and 0,13 (coffee pulp).

The non-parametric analysis of biological efficiency showed that the highest values corresponded to T6 (50.50%) and T5 (49.83%) with significant differences compared to the other treatments. Regarding the production rate, the highest values were achieved with T6 (0.92) and T5 (0.91), with significant differences compared to T1, T2, T4, T7 and T8 (Table 4).

El contenido de proteína (31.57 – 32.16 %) determinado en los carpóforos obtenidos en la presente investigación se encuentra en el rango de 11 – 42 g por 100 g de peso seco reportado para diferentes especies de Pleurotus (Beltrán Delgado et al., 2021) y 17 – 42 g por 100 g de hongos secos para *Pleurotus ostreatus* en residuos sólidos lignocelulósicos (Nieto Juárez et al., 2021). El porcentaje de proteína fue estadísticamente igual en los carpóforos de todos los tratamientos, resultado similar al reportado por (Ríos-Ruiz et al., 2017). Estos investigadores determinaron que en la pulpa de café cultivada con *Auricularia* sp. la eficiencia biológica fue de 30.33 % y la tasa de producción fue de 0.67, valores superiores a los registrados en la cascarilla de arroz que fueron de 19.33 % y 0.43 respectivamente; sin embargo, el contenido de proteína fue similar en ambos substrates (9.01 y 9.30 % respetivamente) (Table 5). The nutritional composition of *Pleurotus ostreatus* carpophores is influenced by the geographical area of substrate (Nieto Juárez et al., 2021), and the protein contained varies according to the composition of the substrate, harvest time, and pili size (Beltrán Delgado et al., 2021).). In this contex, 47.3% of protein was reported in carpophores in coffee grounds crop (Nieto Juárez et al., 2021); 19 % in coffee pulp (Ríos-Ruiz et al., 2017); 13 % in barley straw and 5,9 % in rice husk (Prasad et al., 2023). The nutritional value of *Pleurotus ostreatos* mushrooms is based on the fact that they constitute a source of protein (up to 35 % on dry basic), higher than milk contein (25.2 %) and wheat (13.2 %). It also contains vitamins (B1, B2, B12, C and D), amino acids (niacin, pantothenic acid), polysaccharides, unsaturated fatty acids, phenoles and flavonoids (Beltrán Delgado et al., 2021), therefore, edible mushroom cultivation is a viable alternative for reuse and valorization of waste, reducing environmental impact and providing economics returns to industries (Ritota & Manzi, 2019), statically, protein percentage values did not present significance (Tabla 5).).

**Table 5.** Protein percentage of *Pleurotus ostreatus* basidiocarps grown on different lignocellulosic substrates.

| Tratamientos | Protein* (%) |  |  |  |
| --- | --- | --- | --- | --- |
| | r1 | r2 | r3 | $\bar{X}$ |
| T1: Control rice husk (CA) | 31.68 | 31.70 | 31.84 | 31.74 ± 0.087 |
| T2: Control sugarcane bagasse (BC) | 31.70 | 31.84 | 31.92 | 31.82 ± 0.111 |
| T4: 66,7% CA + 33,3% BC | 31.61 | 33.12 | 31.20 | 31.98 ± 1.011 |
| T5: 50% CA + 50% BC | 31.12 | 31.76 | 31.50 | 31.46 ± 0.322 |
| T6: 25% CA + 75% BC | 31.76 | 31.76 | 32.96 | 32.16 ± 0.693 |
| T7: 66,7% CA + 33,3% C | 31.84 | 32.79 | 31.70 | 32.11 ± 0.593 |
| T8: 50% CA + 50% C | 31.12 | 31.92 | 31.84 | 31.62 ± 0.441 |

\* Kruskal-wallis, p <0,05

## 4. CONCLUSIONS

This study shows that *Pleurotus ostreatus* mushrooms developed in rice husk suplemented with residual biomass from the sugarcane industry and carpophores with a total weight of 83 to 101 g were harvested in rice husk with sugarcane bagasse and 17.33 to 35.67 g in rice husk with sugarcane filter cake compared to 25.67 g in rice husk control. The productivity of *Pleurotus ostreatus* expressed as biological efficiency was 27.66 – 50.50 with rice husk plus sugarcane bagasse and 8.67 – 9.51 with rice husk plus sugarcane filter cake compared to 8.56 in the control rice husk. In turn, the production rate was 0.58 – 0.92; 0.18 – 0.24 y 0.22 respectively, demonstrating that rice husk suplemented with sugarcane bagasse improves performance, production rate, and biological efficiency.

